# Chromatin changes in *Anopheles gambiae* induced by a *Plasmodium falciparum* infection

**DOI:** 10.1101/418442

**Authors:** José L. Ruiz, Rakiswendé S. Yerbanga, Thierry Lefèvre, Jean B. Ouedraogo, Victor G. Corces, Elena Gómez-Díaz

## Abstract

Infection by the human malaria parasite leads to important changes in mosquito phenotypic traits related to vector competence. However, we still lack a clear understanding of the underlying mechanisms and in particular, of the epigenetic basis for these changes. We have examined genome-wide distribution maps of H3K27ac, H3K9ac, H3K9me3 and H3K4me3 by ChIP-seq and the transcriptome by RNA-seq, of midguts from *Anopheles gambiae* mosquitoes infected with natural isolates of the human malaria parasite *Plasmodium falciparum* in Burkina Faso. We report 15,916 regions containing differential histone modification enrichment, of which 8,339 locate at promoters and/or intersect with genes. The functional annotation of these regions allowed us to identify infection responsive genes showing differential enrichment in various histone modifications, such as CLIP proteases, anti-microbial peptides encoding genes, and genes related to melanization responses and the complement system. Further, the motif analysis of regions differentially enriched in various histone modifications predicts binding sites that might be involved in the cis-regulation of these regions such as Deaf1, Pangolin and Dorsal transcription factors (TFs). Some of these TFs are known to regulate immunity gene expression in *Drosophila* and are involved in the Notch and JAK/STAT signaling pathways. The analysis of malaria infection-induced chromatin changes in mosquitoes is important not only to identify regulatory elements and genes underlying mosquito responses to a *P. falciparum* infection but also for possible applications to the genetic manipulation of mosquitoes and to other mosquito-borne systems.

## INTRODUCTION

Mosquitoes are the most medically important arthropod vectors of disease. Among mosquito-borne diseases, malaria, caused by protozoan parasites of the genus *Plasmodium*, is the deadliest. The human malaria parasite *P. falciparum* is the most prevalent agent in Africa and it is responsible for at least 200 million acute cases worldwide and around half a million deaths each year [1]. It is transmitted by *Anopheles* mosquitoes, *Anopheles gambiae* being the main vector. At the molecular level, *P. falciparum* infection induces drastic and rapid changes in gene expression in mosquito tissues that relate to functions involved in immunity, development, physiology, and reproduction [2]. The immune response during infection is the best characterized of these pathways, and mostly occurs in the midgut of infected mosquitoes. This involves, among others, the activation of genes involved in epithelial nitration responses, melanization and the complement-like system [3-5]. Furthermore, infection in mosquitoes impacts epidemiologically important life-history traits such as vector competence, i.e. the ability of the mosquito to acquire, maintain, and transmit the parasite, and prime subsequent infections [6-8]. There is substantial variability in the responses of the mosquito to *P. falciparum* infection that depends on both genetic and environmental contexts [9, 10]. However, the mechanisms that regulate phenotypic responses to infection in mosquitoes, and that might mediate memory of the malaria infection-phenotype, are little understood.

Chromatin-associated processes participate in the regulation of gene expression during development as well as in the differentiated tissues of the adult organism. These processes are sensitive to environmental stimuli, they are more or less transitory and can potentially be inherited, allowing living organisms and individual cells to continuously integrate internal and external inputs and to mediate responses through gene regulation [11, 12]. Amongst these, post-translational modifications of histones (hPTMs) impact the structure and/or function of chromatin, with different histone marks yielding distinct functional consequences [13]. For example, in *Drosophila*, as in other organisms, H3K27ac, H3K9ac and H3K4me3 are linked to gene activation and localize at promoters, whereas H3K9me3 is associated with silencing and occupies broader regions [14]. Various combinations of active and repressive histone modifications define chromatin states that are linked to gene function [15-17]. These modifications can remain during cell division, leaving a record of gene activity, i.e. epigenetic memory, that affects or primes the transcriptional response later in life [12, 18-24]. In mosquitoes, despite their relevance to human health, there is very little knowledge of chromatin regulation and its link to mosquito immunity, physiology and behavior [25]. In a previous study, we characterized genome-wide occupancy patterns of various histone modifications and established a link between hPTMs and gene expression profiles in midguts and salivary glands of the human malaria vector *An. gambiae* [26]. In addition, recent reports revealed the role of various transcription factors such as REL2, lola and Deaf1, in *An. gambiae* immune defenses [27, 28]. These findings have set the stage for additional studies aimed at understanding how the chromatin landscape is altered in *P. falciparum-*infected mosquito tissues and what are the molecular players involved in these malaria-induced responses.

Available data on the phenotypic and transcriptional responses of mosquitoes to infection by *Plasmodium* is built on the use of mosquito-parasite combinations that in most cases differ from those found in nature. In these studies, infections often take place under standard laboratory conditions, using laboratory-adapted parasite clones and commercially available laboratory mosquito strains. Such experiments are useful to distinguish the contribution of parasite and mosquito genetic factors and also the influence of various environmental variables on the infection output, but they may not reflect the complexity of interactions that take place in natural conditions [29]. In this context, field-based studies are critical as they offer the advantage of allowing a more realistic picture of the molecular interactions in the context of natural transmission.

In this study, we aim to provide a comprehensive understanding of the chromatin and the transcriptional responses induced by *P. falciparum* infection of *An. gambiae* in the conditions found in a malaria endemic area in Africa. For this purpose, we compared genome-wide maps of histone modification profiles in infected and control mosquito tissues to identify chromatin state transitions associated with infection, and examined the expression pattern of genes that annotate to regions containing differential histone modifications using the natural association between the mosquito *An. gambiae* and natural field isolates of the human malaria parasite *P. falciparum* in Burkina Faso. Motif enrichment analysis at these regulatory regions allows us to predict the binding sites of several transcription factors, some of which have been shown to be involved in mosquito immune responses by previous studies.

## RESULTS

### Chromatin states in malaria-infected mosquitoes

*An. gambiae* were fed with blood obtained from malaria-infected human volunteers in Burkina Faso. Midguts from infected and uninfected (blood fed control) female mosquitoes were dissected and pooled separately for each condition. On the pooled samples we carried out RNA-seq and ChIP-seq using antibodies to several histone modifications: H3K9ac, H3K27ac, H3K4me3 and H3K9me3. Regions enriched in these histone modifications identified by MACS2 [30] are listed in Table S2. We found similar numbers of peaks in the infected and uninfected mosquitoes and a high correlation in the ChIP-seq profiles (Figure S1A), indicating that histone modification profiles are comparable between the two conditions. Based on this result, we focus particularly on characterizing functional chromatin states in the infected, and the results for the uninfected are in the supplemental material (Figures S1 and Figure S2).

ChIP-seq peaks annotated to genomic features are shown in Figure 1A. The analysis of ChIP-seq peaks with respect to genomic features shows that upstream regions significantly enriched in histone modifications are mostly located in a 2 Kb window from the translation start codon (ATG) of *An. gambiae* genes, with lower enrichment at distances greater than 2 Kb (Figure 1B). Based on these findings, we annotated peaks to the promoters of nearby genes when located less than 2 Kb upstream, and used this annotation for all subsequent analysis. Results show that most ChIP-seq peaks correspond to H3K9me3 marked regions (37,343) followed by H3K27ac (35,217), H3K4me3 (19,945) and H3K9ac (6,131) (Table S2).

**FIGURE 1.**
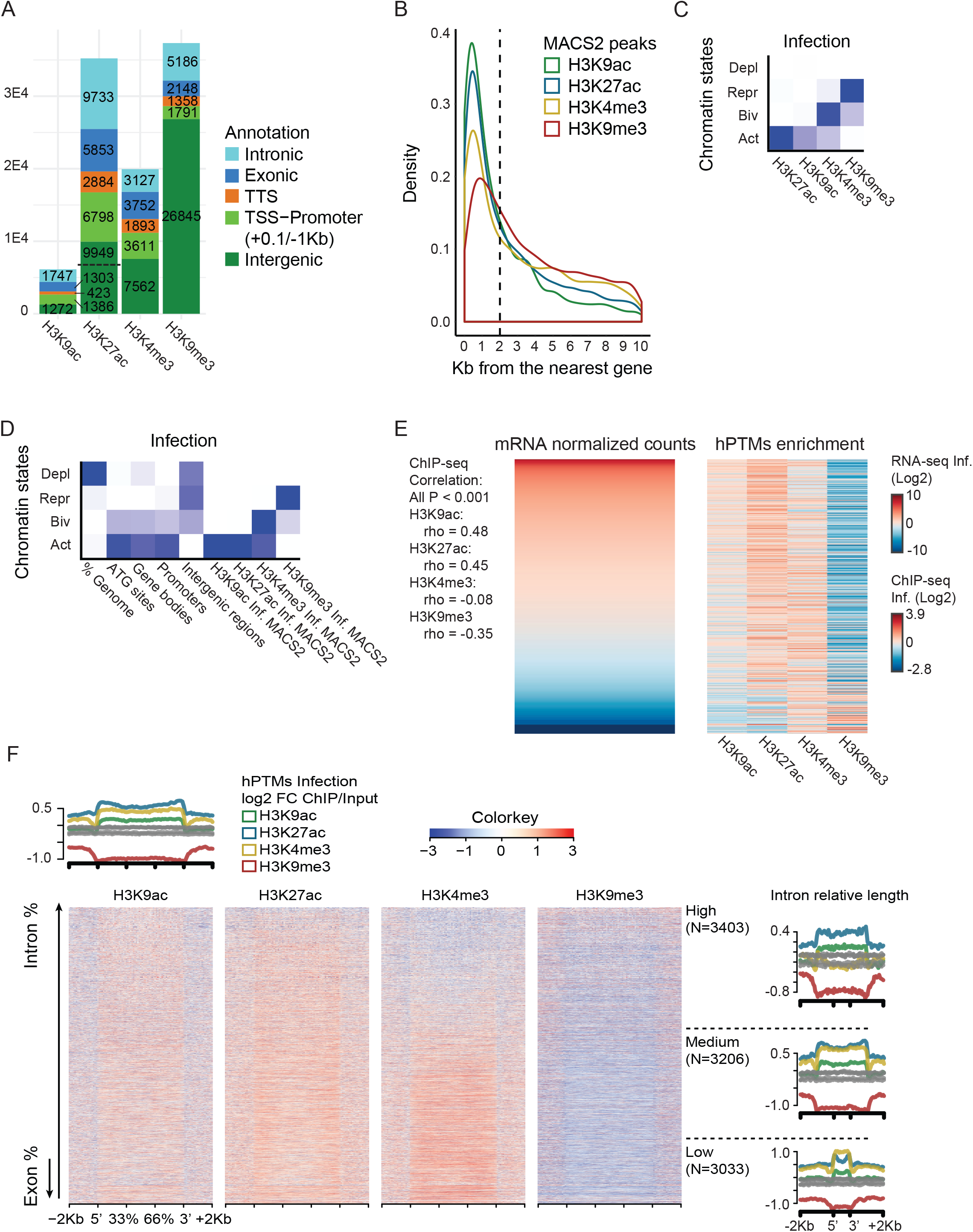
Histone modifications profiles and gene expression regulation in *P. falciparum*-infected mosquitoes. A. Annotation of MACS2 ChIP-seq peaks for each histone modification to genomic features: TSS-Promoters, TTSs, Intergenic, Intron and Exon regions. B. Density plot showing the position (Kb upstream) of MACS2 peaks for each histone modification with respect to the ATG protein initiation codon of the nearest downstream gene. C. Heatmap of emission parameters from ChromHMM analysis using a four chromatin-states model based on histone modification enrichment patterns in the infected condition. The predicted states are: Deplet (depleted, low levels of all hPTMs), Repr (repressive, H3K9me3 enrichment), Biv (bivalent, H3K4me3/H3K9me3 enrichment) and Act (active, H3K27ac/H3K9ac/H3K4me3 enrichment). Darker blue indicates higher enrichment of a particular histone modification. D. Heatmap showing the overlap of various genomic features, including MACS2 peaks located in promoters (2 Kb from the ATG) or gene bodies in the infection condition, with the predicted chromatin states. Darker blue in the first column indicates higher percentage of the genome overlapped by a given state. For other columns it indicates the likelihood of finding a particular chromatin state in each genomic feature compared to what it would be expected by chance. E. Heatmaps showing mRNA levels (left) and histone modification enrichment profiles (right) of genes displaying a MACS2 peak in the promoter or the gene body. Data corresponds to the infected condition. Genes are ordered by mRNA levels. ChIP-seq enrichment at the promoters and the gene bodies is normalized (RPKM) and input-corrected. Data is log2-scaled and mean-centered. Spearman rank correlation coefficient (rho) and corresponding P-value are shown for the association between each histone modification enrichment levels and mRNA levels. F. Heatmaps showing histone modification enrichment profiles at high and medium expressed genes in the infection condition. Genes are ordered by the percentage of the body containing introns and exons. Average profile plots show density of normalized (RPKM) and input corrected ChIP-seq reads for each histone modification at high and medium expressed genes (top) and at those genes classified by the percentage of the gene body containing introns (right).

We used ChromHMM [31] to segment the genome into 4 distinct chromatin states based on relative enrichment levels of H3K9ac, H3K27ac, H3K4me3 and H3K9me3, that we named as follows: hPTM depleted, i.e. low levels of enrichment for all hPTMs, active (H3K9ac/H3K27ac and H3K4me3), repressed (H3K9me3), and bivalent (H3K4me3/H3K9me3) (Figure 1C, Figure S1B). As expected, most of the genome is in a depleted state, whereas gene bodies and promoters display enrichment in active state (Figure 1D, Figure S1C). Following this categorization, we investigated the association between various combinations of active and repressive histone modifications at promoters and gene bodies, and gene expression by RNA-seq. As expected, active chromatin is associated with expressed genes, whereas chromatin marked with H3K9me3 associates with silent or low-expressed genes. Genes showing a bivalent enrichment pattern (H3K4me3 and H3K9me3) are generally expressed at low levels (Figure 1E, Figure S2A). This is corroborated by a positive and statistically significant correlation between levels of enrichment in active histone modifications (H3K9ac, H3K27ac, H3K4me3) and mRNA levels of ChIP-seq peaks annotated genes, whereas the correlation is negative for H3K9me3 (Figure 1E, Figure S2A). Following what has been described for *Drosophila* [32], we classified genes according to the gene structure, i.e. length of the coding region and length of intronic segments. We observed that expressed genes with long introns and relatively short coding regions show broad H3K27ac and H3K9ac domains. In contrast, expressed genes with more uniformly distributed coding regions show a more localized H3K27ac/H3K9ac enrichment through the gene body and higher enrichment in H3K4me3 (Figure 1F, Figure S2B).

### Regions differentially enriched in histone modifications associated with P. falciparum infection

We find a considerable overlap in histone modification profiles between infected and uninfected mosquitoes, with more than 30,000 common ChIP-seq peaks between the two conditions. However, a portion of the peaks appears to be condition-specific (8,234 in infected and 14,138 in the uninfected) (Figure S2C). Based on this observation, we used the diffReps software [33] to further investigate localized changes in hPTMs enrichment that might occur in response to *P. falciparum* infection. We identified 15,916 regions containing significantly different levels of ChIP-seq signal (P-value < 10E-4) for all four histone modifications. The number of diffReps regions was similar for H3K9ac (2,396) and H3K4me3 (2,837), whereas H3K27ac and H3K9me3 displayed a larger number of differential regions (4,810 and 5,873, respectively) (Table S3). Regions of differential active histone modifications between infected and control mosquitoes were primarily distributed near genes, upstream and downstream, or in introns. But they also occupied distal intergenic sites, particularly in the case of H3K9me3 (Figure 2A-B). We applied a series of filtering thresholds (see Methods) to these differential regions to obtain a high confidence set that we classified according to chromatin state transitions (ChromHMM) (Figure 2C). In the majority of cases, the diffReps changes involved an enrichment or depletion in a certain histone modification without a chromatin state change, but we also reported chromatin state shifts between conditions: regions that were active upon infection or regions that were marked with active chromatin marks in control mosquito tissues and changed to depleted in the infected. There was also a considerable proportion of regions that switched between the depleted and H3K9me3 enriched states (Figure 2C).

**FIGURE 2.**
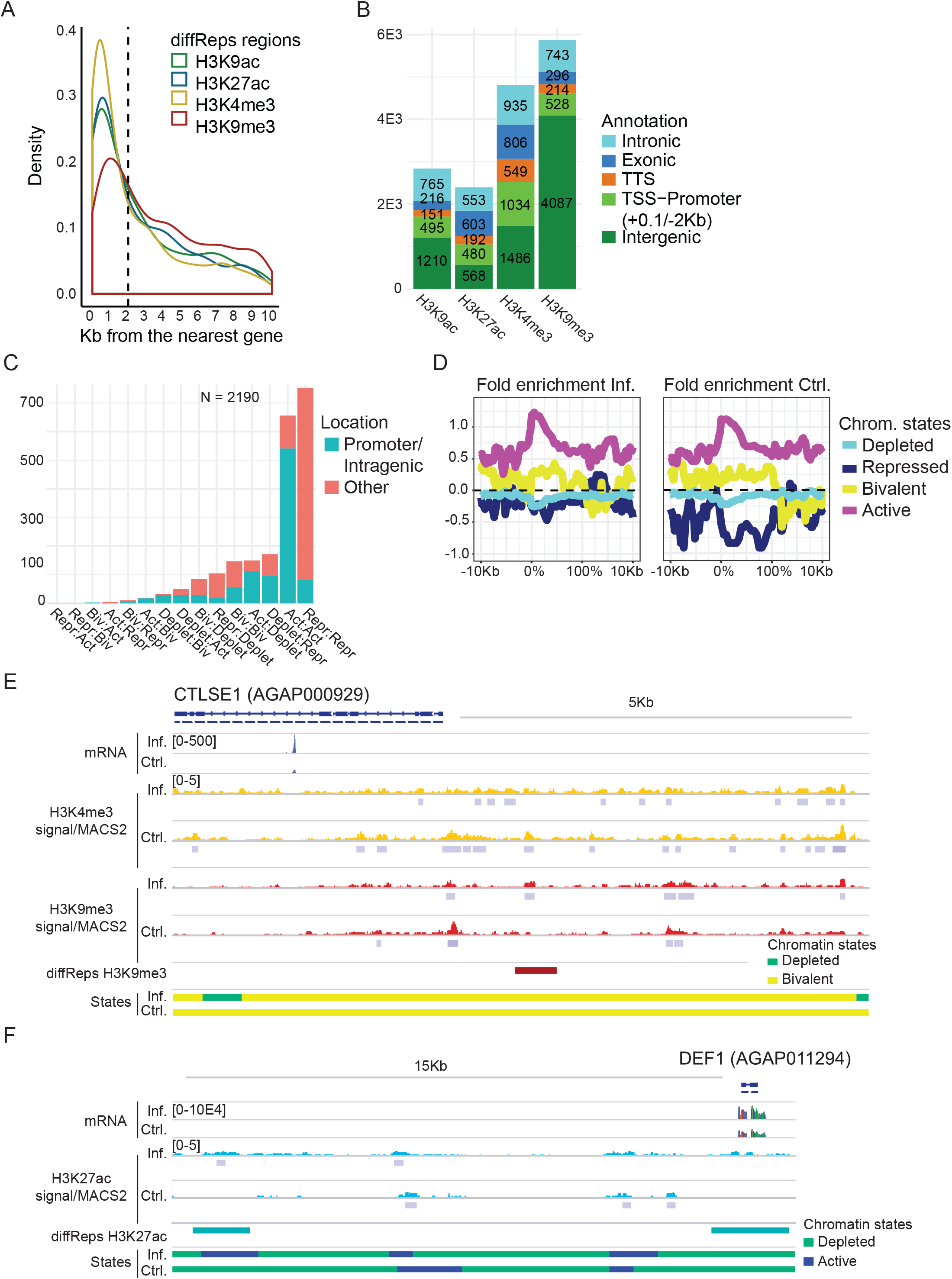
Changes in histone modification enrichment in response to infection. A. Density plot showing the position (Kb upstream) of differential diffReps regions for each histone modification with respect to the ATG initiation codon of the nearest downstream gene. B. Annotation of diffReps regions for each histone modification to genomic features: TSS-Promoters, TTSs, Intergenic, Intron and Exon regions. diffReps regions located -2 Kb/+0.1 Kb from the ATG are annotated to TSS-Promoter regions. C. Barplot showing the number and location of high confidence diffReps regions and the chromatin state transitions between conditions associated with the peak region. D. Profile plots showing predicted chromatin states in infection (left) and control (right) conditions at genes encoding for immune-related factors [1]. The graphs represent chromatin state fold enrichment (log(observed/expected)) with respect to the scaled gene bodies +- 10 Kb. E-F. Histone enrichment profiles in the regions containing the CTLSE1 (AGAP000929) and DEF1 (AGAP011294) encoding genes. Tracks show normalized/input-corrected ChIP-seq signals and RNA-seq mapped read counts for each condition. The location of diffReps regions, MACS2 peaks, and predicted chromatin states for each condition are included. All tracks are shown at equal scale.

The gene ontology analysis shows that diffReps-annotated genes appeared significantly enriched in GO terms related to development, transcription regulation and metabolism (Table S3). In addition to this analysis, we also looked for coincidences between the diffReps-annotated genes and genes that have been reported to be involved in the immune response to infection [34, 35]. Among diffReps-annotated genes, we found 133 genes encoding proteins involved in the immune response (26 considering the high confidence set of diffReps peaks). Genes differing between conditions in their histone modification profiles encode proteins involved in apoptosis (IAP3 and IAP7), Clip-domain serine proteases and serine protease inhibitors (CLIPC, CLIPE, SRPN10 and SRPN4), C-type lectins (CTLs, Figure 2E), antimicrobial peptides (DEF1, Figure 2F), Scavenger Receptor (SRCR domain) with Lysyl Oxidase domain (SCRAL1), and components of Toll, NF, and peptidoglycan recognition protein LC/immune deficiency (PGRP-LC/IMD) signaling pathways, among others (Table S3, Figure S3). Most genes involved in immune responses were enriched in the same chromatin states in both conditions but displayed a change in the relative abundance of active H3K4me3/K9ac/K27ac, and/or the repressive H3K9me3 modification (Figure 2D, Table S3). There were 9 genes within the high-confidence set of diffReps-annotated genes that displayed differential chromatin state between conditions like the long caspase CASPL2 encoding gene and the fibrinogen-related protein encoding gene (Table S3, Figure S3B-C).

### Complex relationship between chromatin and gene expression changes

In order to investigate the functional significance of the regions differentially enriched in various histone modifications, we selected high confidence differential hPTMs regions that overlap gene bodies or are located up to 2 Kb upstream of genes. This analysis resulted in the identification of 1,208 genes for all four hPTMs. We then applied a soft clustering approach using Mfuzz [36] to the -2 Kb-gene region. This analysis allowed us to group diffReps-annotated genes based on unique profiles of hPTM enrichment (Figure S4) and to examine the correlation between changes in the histone modification patterns and the expression status. We found that genes differentially enriched in a condition in active histone marks (H3K27ac, H3K9ac and H3K4me3) tended to be display high expression in that condition, whereas those that were marked with repressive H3K9me3 or bivalent H3K4me3/H3K9me3 tended to be display low expression (Figure 3A, Figure S5A, Table S4). However, when comparing the ratio of histone enrichment versus the ratio of gene expression values between infected and control mosquitoes, the correlation coefficient was low and non-significant, meaning that the infection condition influenced to a different extent chromatin and gene expression patterns. This was clearly shown when examining the ratio of enrichment of various hPTMs in the infected relative to the control, for infected expressed genes (left panel) or control expressed genes (right panel) (Figure 3B). The ratio was in the same direction, above or below 1 for infected and control Log2 values respectively, only in the case of the H3K9me3 (Figure 3B). Nevertheless, this integrative analysis identified 278 genes in which the differential active or repressive histone modification enrichment coincided with a shift in the gene expression pattern between the infected and control condition (Table S4, Figure 3C, Figure S3B).

**FIGURE 3.**
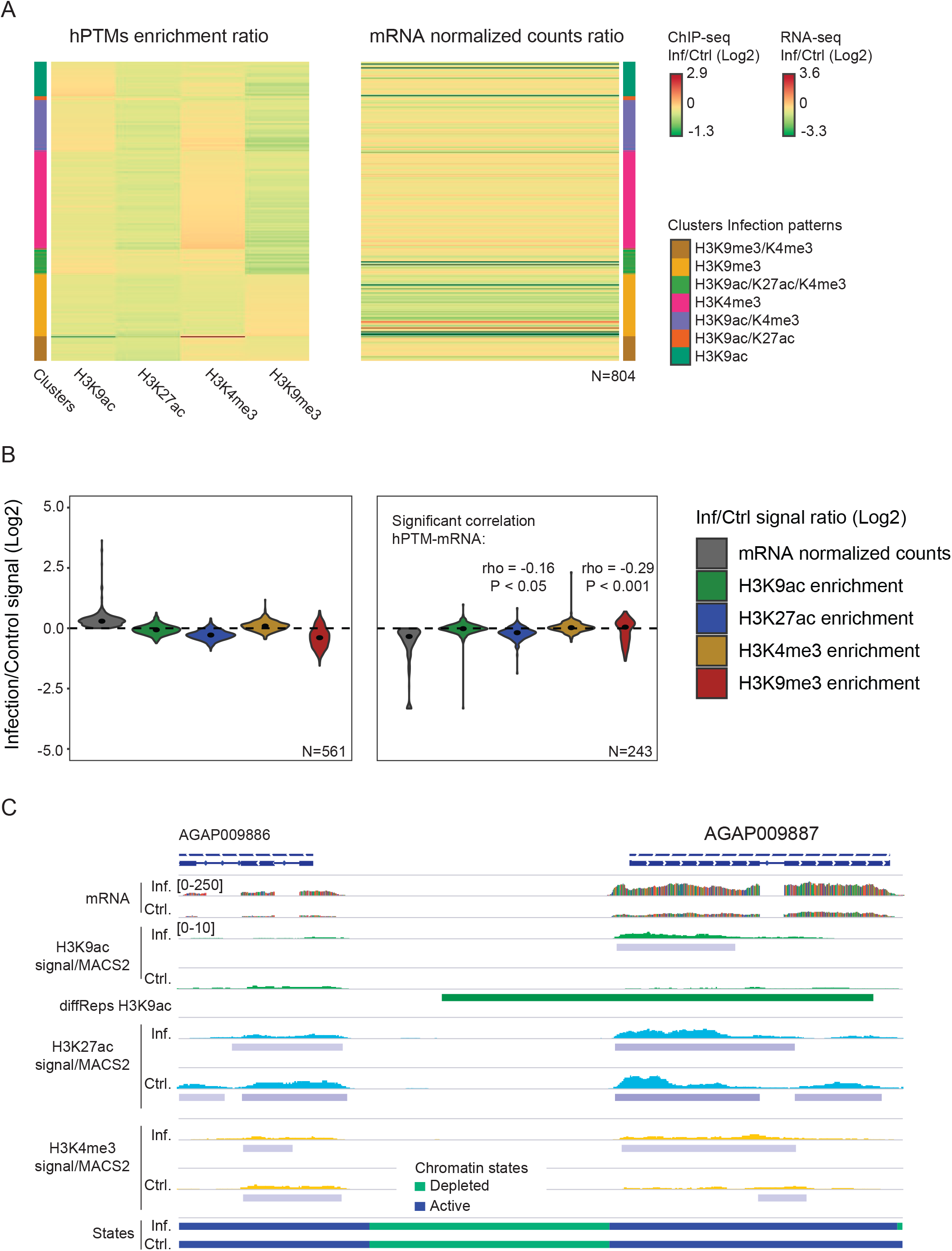
Association between histone modification differential enrichment and changes in gene expression. A. Heatmaps showing clusters of genes (-2 Kb) grouped by unique histone modification profiles identified in the soft clustering analysis (left) and corresponding changes in mRNA levels (right). ChIP-seq enrichment at the promoters and gene bodies is normalized (RPKM) and input-corrected. The signal corresponds to the ratio of ChIP- seq and mRNA levels in the infected versus the control condition. Data is log2-scaled and mean-centered. B. Ratio of gene expression and histone modification enrichment between infected and control conditions for Mfuzz gene clusters more highly expressed in infected (left) and control (right) conditions. Data is the log2-scaled ratio between the infected and the control as in (A). Spearman rank correlation coefficient (rho) and corresponding P-value are shown for significant correlations between histone modification enrichment and mRNA levels. C. Histone enrichment profiles in the region containing the AGAP009887 coding gene. Tracks show normalized/input-corrected ChIP-seq signals and RNA-seq mapped reads counts for each condition. The location of diffReps regions, MACS2 peaks, and predicted chromatin states for each condition are included. All tracks are shown at equal scale.

In addition to the analysis of the diffReps-annotated genes, we also examined differentially expressed genes and looked for differential chromatin marks between conditions. The DESeq2 analysis on the RNA-seq data revealed 713 significantly different expressed transcripts (P-value < 0.05, 184 up-regulated genes and 529 down-regulated genes in the infected vs. the control condition) (Table S5). We found 105 differential expressed genes that contain a diffReps region annotated to the promoter or the gene body. Of those, there were 72 in which the direction of the change (active or repressive histone marks) agreed with the functional prediction (up or down-regulation) (Table S5). Same as above, the switch in the expression status was generally linked to changes in the H3K9me3 enrichment levels (Figure S6A). Examples of diffReps/DESeq2 genes were IAP7 (AGAP007293) and Argonaute 4 (AGAP011717). IAP7 was upregulated in infection and H3K9me3 was depleted in this condition compared to the control (Figure 4A), while AGAP011717 was expressed at higher levels in the uninfected and this was associated to a gain in active histone modifications, mainly H3K27ac (Figure 4B).

**FIGURE 4.**
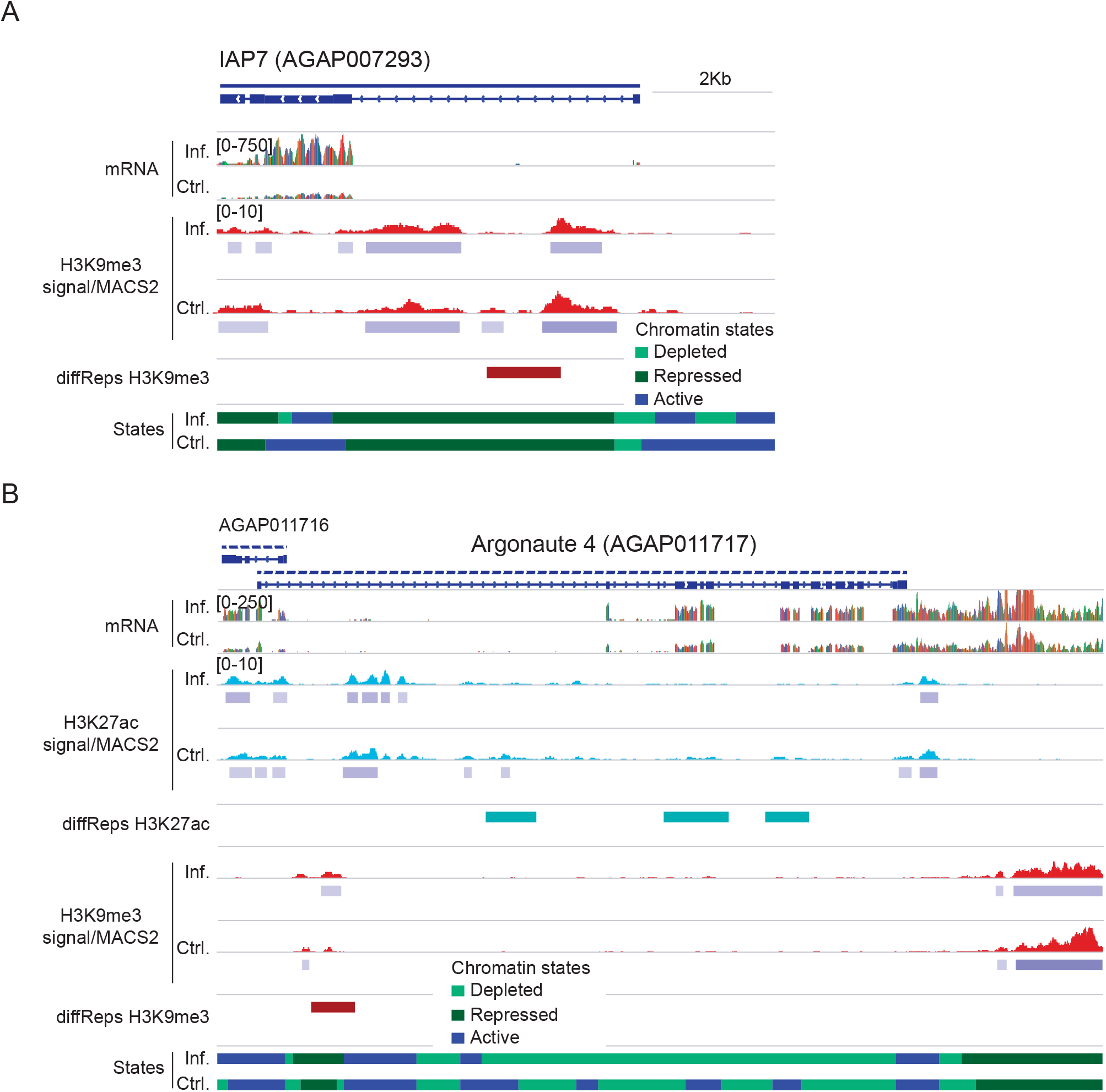
Significant differential gene expression and association with histone modification differential enrichment. A-B. Histone enrichment profiles in the regions containing the IAP7 (AGAP007293) and Argonaute 4 (AGAP011717) encoding genes. Tracks show normalized/input-corrected ChIP-seq signals and RNA-seq mapped reads counts for each condition. The location of diffReps regions, MACS2 peaks, and predicted chromatin states for each condition are included. All tracks are shown at equal scale.

### Motif analysis of regions differentially enriched in histone modifications identifies transcription factor binding sites involved in transcriptional responses to infection

We conducted DNA-binding motif enrichment analysis on the set of ChIP-seq high confidence diffReps regions that coincided with MACS2 peaks of significant enrichment, which includes 2,018 peaks for all four histone modification marks (Table S3). Such an approach is useful for histone modification peaks because it allows the discovery of unanticipated sequence motifs associated with specific histone marks, like transcription factor binding sites. This analysis revealed multiple novel motifs that are significantly enriched in sequences containing H3K9ac, H3K27ac, H3K4me3, and H3K9me3 differentially enriched peaks. Table 1 lists novel motifs identified by HOMER on the set of diffReps peaks for each histone modification and their similarities with known TF binding sites previously described in *Drosophila*. We found that the binding sites predicted at ChIP-seq peaks showed similarities with those of transcription factors such as Deaf1, pangolin (pan) and Dorsal (Dl), linked to immunity gene expression regulation in *Drosophila*. We also found binding sites for several transcription factors associated with developmental functions such as Caudal (Cad), and reproduction like Eip74EF (Table 1).

**TABLE 1.**
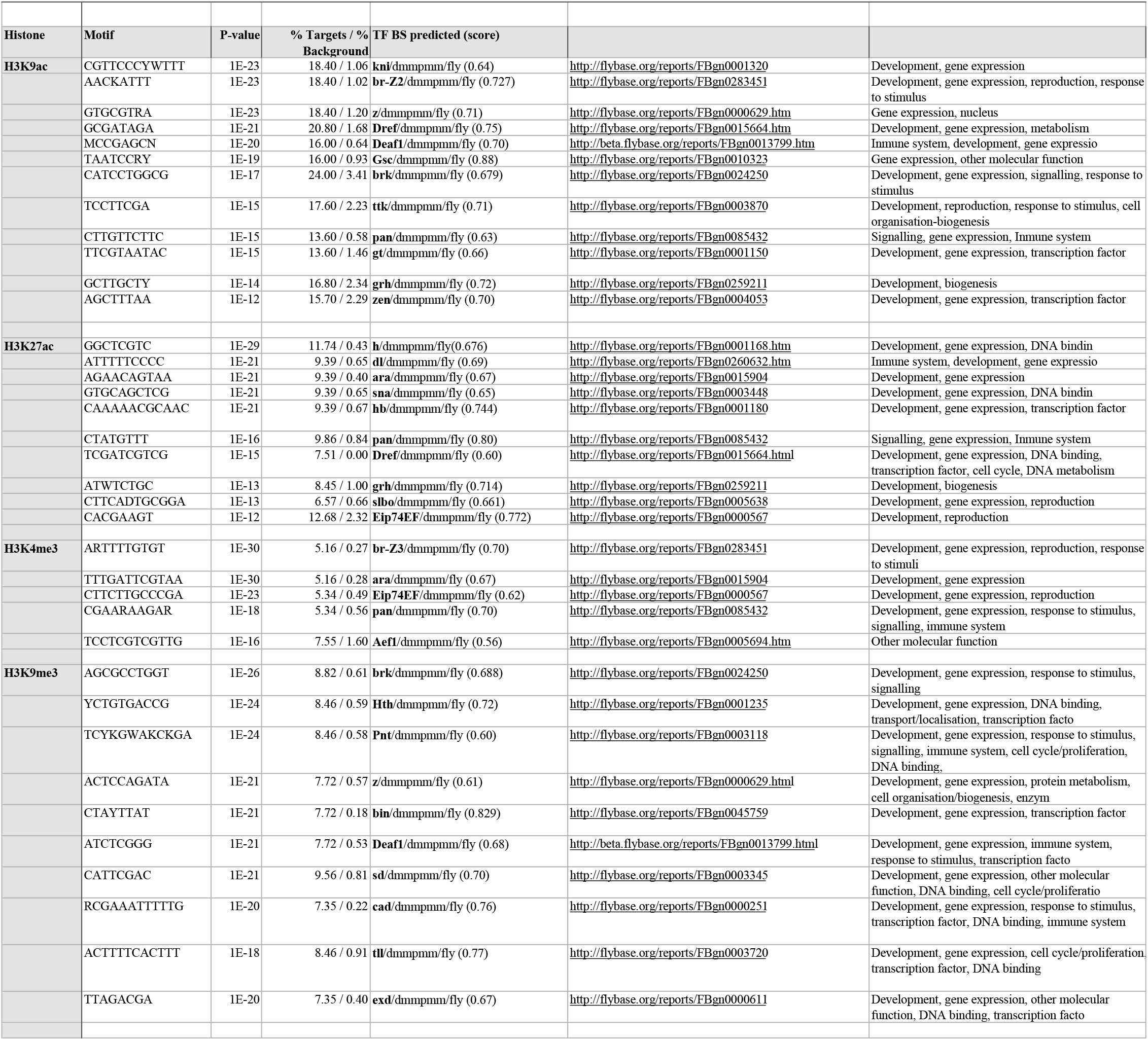
List of consensus motifs corresponding to known transcription factors significantly enriched in the set of differential histone modified regions that are associated to a *P. falciparum* infection.

As a validation of our strategy to predict regulatory sites and TF binding, we reported the overlap between FAIRE-seq peaks determined in *An. gambiae* hemocytes in a previous study, with histone modifications ChIP-seq peaks in malaria-infected tissues. Even if the experimental conditions and cell types were not comparable, we found 9,136 MACS2 ChIP-seq peaks and 2,690 diffReps regions (10% and 17% of the total, respectively) that intersected with FAIRE-seq peaks.

These results collectively suggest that some of the transcription factors reported in our study are probably involved in chromatin remodeling processes and the regulation of transcription of the genes that are elicited in mosquitoes in response to a *P.falciparum* infection.

## DISCUSSION

Human malaria is a mosquito-borne disease responsible for around half-million deaths per year, *An. gambiae* being the main disease vector in Africa [1]. An infection by *P. falciparum* alters the phenotype and vector competence of mosquitoes with consequences for transmission and malaria epidemiology. However, the molecular players that regulate malaria infection-triggered responses are still poorly known. A considerable amount of work exists on the genomic basis of mosquito resistance to infection (reviewed by [37]), but there is a dearth of epigenomic studies on the relationship between chromatin and gene expression regulation in mosquitoes. In a previous study, we characterised for the first time genome-wide profiles of various histone modifications in *An. gambiae* and compared these patterns with chromatin maps published for *Drosophila* [26]. Here, we go a step further and performed an integrative analysis of ChIP-seq and RNA-seq data in the context of malaria infection in the natural conditions of transmission of the disease.

We report various chromatin states with links to functional gene expression in the study of global patterns of histone modifications in infected mosquitoes. H3K9ac/H3K4me3/H3K27ac histone acetylation marks are associated with the promoters of active genes and repressive H3K9me3 is associated with silent genes. In agreement with previous studies in *Drosophila*, our results in mosquitoes show that gene structure is related to differences in the distribution of H3K27ac and H3K4me3 enrichment in the gene body, being the most common difference in the chromatin state of expressed genes that contain long introns relative to exons. We also report bivalent chromatin domains, that have both repressive H3K9me3 and activating H3K4me3 histone marks in the same region. This pattern is typical of transcription factor genes that are expressed at low levels, and have been also found associated with genes involved in development as well as gene imprinting [38, 39]. Despite the fact that the majority of peaks are present at promoters or gene bodies, we report a considerable number of ChIP-seq peaks (~40%) located more than 2 Kb from the nearest gene. It has been described that H3K27ac associates with active enhancers in *Drosophila* and other model organisms [40, 41]. In mosquitoes, we identified a number of distal and intragenic H3K27ac-enriched regions that could correspond to enhancer-like sites. Future ChIP-seq studies on enhancer-specific marks combined with chromatin accessibility mapping are required to confirm the presence of enhancer-like elements in mosquitoes.

Once we showed a relationship between chromatin composition and function, our main purpose was to identify malaria-induced chromatin changes in mosquitoes. We could identify 15,916 histone modified regions (2,564 when considering high confidence peaks) that appeared differentially methylated or acetylated upon infection. Various chromatin states -active, repressive and bivalent chromatin - were identified in the set of regions marked by differential levels of histone modifications. We also observed that differential regions generally display enrichments or depletions of individual marks at specific gene segments, but maintaining the same chromatin state. Importantly, there were 107 promoter or intragenic regions and 133 more distal sites (>2 Kb), that correspond to genes involved in different pathways of mosquito innate immunity either as activators or inhibitors of various responses (see [42, 43] for reviews). These include apoptosis, Clip-domain serine proteases and serine protease inhibitors, antimicrobial peptides, enzymes that catalyze generation and detoxification of reactive oxygen species, and components of Toll, NF and IMD immune signaling pathways.

The integration of regions differentially marked by histone modifications with expression levels of annotated genes resulted in the identification of 278 malaria-responsive genes. These genes show local differences in the enrichment of specific active/repressive histone marks that correlate with gene expression changes in the same direction. However, this is not the general rule and most of the differential regions by ChIP-seq do not display noticeable differences in gene expression, and the other way around for differentially expressed genes. It might be that there is a threshold enrichment level that is necessary to activate transcription. It is also unexpected that comparison of infected with control tissues only identifies a few of these malaria responsive genes corresponding to factors involved in the immune response. A possible explanation is that the majority of the immune response factors that have been described belong to the early innate immunity response that takes place between 2-24h post-infection, at the ookinete stage, where the most part of the parasite recognition and killing occur [44, 45]. The samples analyzed in this study correspond to the oocyst stage, approximately six and seven days after an infective blood feeding, and the immune factors playing a role at this stage still remain poorly characterized [46].

Cis-regulatory elements are implicated in the control gene expression because they contain specific DNA sequences that are binding sites for transcription factors and other chromatin remodelers and often appear enriched in certain histone modifications. These elements are well mapped in *Drosophila* [47], but very few have been identified in mosquitoes [27, 48]. The analysis of differential ChIP-seq peaks located at promoters or gene bodies identified significant enrichment in binding sites that match consensus sequences of TFs previously described in *Drosophila*, including transcription factor deformed epidermal autoregulatory factor-1 (DEAF-1) [49]. Indeed, DEAF-1 is an important regulator of *Drosophila* immunity that could induce genes encoding antimicrobial peptides [50]. Another example that we report in this study is Dorsal (Dl), a TF that functions downstream of the Toll pathway [49]. Finally, as a validation of our motif analysis, we find that a portion of the differential ChIP-peaks of histone modifications matches FAIRE-seq peaks described by a previous study on mosquito hemocytes [27]. In this study, authors reported that FAIRE sequences were enriched in binding sites for DEAF-1, which is one of the top motifs reported in the present study. New approaches to profiling chromatin accessibility such as ATAC-seq will be useful to further characterize cis-regulatory sequences and TF binding *in vivo*.

In summary, this study charts genomic landscapes of various active and repressive histone modifications in malaria-infected mosquitoes and integrates these profiles with RNA-seq data to quantify gene expression. Using this approach, we have identified malaria-responsive genes that display changes in the abundance of specific histone modifications. However, the relationship between chromatin and gene expression changes at differential regions is complex, and only a subset of genes shows correlated patterns that agree with the predicted function. Further research to identify regulatory sequences associated with these changes and the transcription factors with which they associate could provide new molecules and targets for vector control.

## METHODS

### Mosquito rearing and experimental infections

Three- to five-day-old female *An. gambiae* mosquitoes were sourced from an outbred colony established in 2008 and repeatedly replenished with F1 from wild-caught mosquito females collected in Soumousso near Bobo-Dioulasso, south-western Burkina Faso (West Africa). Mosquitoes were maintained under standard insectary conditions (27 ± 2°C, 70 ± 5% relative humidity, 12:12 LD). Two independent experimental infections, biological replicates, were carried out by membrane blood feeding in the laboratory as described previously [51-54]. Briefly, females were fed through membranes on gametocyte-infected blood from malaria patients. Venous blood was collected and the serum was replaced by a non-immune AB serum to avoid transmission of human blocking factors. Dissection of mosquito midguts and salivary glands was performed in situ on adult females at 7 days post-blood meal. To determine infection levels, mosquito guts were dissected 7 days after blood feeding and were stained with 2% Mercurochrome before microscopic examination. Tissues were maintained in ice-cold Schneider’s insect culture medium (Sigma-Aldrich) and fresh tissues were immediately processed for chromatin and RNA analyses.

### RNA isolation, library preparation and sequencing

We prepared RNA-seq libraries from RNA isolated from two biological replicates of uninfected and infected midgut samples. Total RNA was extracted from fresh mosquito tissues (~25 midguts and ~ 50 salivary glands) using the mirVana™ RNA Isolation Kit (Ambion®) according to the manufacturer protocol and used for mRNA library preparation. RNA concentration was quantified using a Qubit^®^ 2.0 Fluorometer, and RNA integrity was determined with an Agilent 2100 Bioanalyzer. Illumina libraries were prepared and sequenced at the HudsonAlpha Institute for Biotechnology, using an Illumina HiSeq2000 sequencer, standard directional RNA-seq library construction, 50 bp paired end reads with ribosomal reduction (RiboMinus™ Eukaryote Kit, Ambion^®^).

### RNA-seq analysis

We mapped RNA paired directional reads to *An. gambiae* PEST strain genome version 4.3 publicly available at Vector Base (https://www.vectorbase.org/) using TopHat v2.0.13 [55]. We aligned reads using the option of library type set as first-strand for directional RNA-seq. We used SAMtools v.1.6 (http://samtools.sourceforge.net) for SAM and BAM file manipulation and conversion. We performed quality control analysis using QualiMap v2.2.1 [56]. Statistics of the RNA-seq analysis for each condition and replicate are shown in Table S1.

Quantification and differential gene expression analysis was conducted using HTSeq/DESeq2 packages [57, 58]. To count reads, HTSeq configuration parameters were set for a strand-specific assay to separate between sense and antisense transcripts. We used the matrix of raw read counts as input for DESeq2 R package, which performs library normalization and uses negative binomial generalized linear models to identify differentially expressed genes. The design included condition as main factor and infection as covariable to control for inter-experiment variability. In this analysis, genes were considered significantly differentially expressed if the P-value was below 0.05.

Sets of differentially expressed genes between conditions, infected and control, were annotated based on the Gene Ontology (GO) and Kyoto Encyclopedia of Genes and Genomes (KEGG) using DAVID [59, 60].

### Chromatin Immunoprecipitation

Chromatin immunoprecipitation in mosquito tissues was performed as previously described [26]. Antibodies to histone modifications used in this study were anti-H3K9ac (Millipore #07-352), anti-H3K4me3 (Abcam ab8580), anti-H3K27ac (Abcam ab4729), and anti-H3K9me3 (Abcam ab8898). ChIP-seq libraries were prepared following the procedure described by Bowman et al. [61], and using the HiFi Kapa Sybr library preparation kit (KapaBiosystems). To obtain the quantity needed to perform ChIP-seq, the two samples for which we have RNA-seq expression data were pooled, resulting in one biological replicate for infected mosquito midguts and one biological replicate of un-infected tissues. ChIP-seq libraries were sequenced at the HudsonAlpha Institute for Biotechnology using an Illumina HiSeq2000 sequencer.

### ChIP-seq data analysis

We performed quality control of Illumina reads using QualiMap v2.2.1 [56] (Table S1). Correlation analysis was performed using deepTools2 (v2.5.0) [62]. We reported Spearman *rho* correlation coefficients between each pair of histone modification datasets.

We mapped reads for various histone modifications and the input to *An. gambiae* PEST strain genome version 4.3 (https://www.vectorbase.org/) using Bowtie v2.2.9 [63] with default parameters except for –no-mixed. Reads were trimmed 5 bases from each read 5’ end (–trim5 5). Mapped reads were then sorted and deduplicated using SAMtools v.1.6. We applied a quality threshold of 10 in MAPQ score. All libraries were downsampled (SAMtools) to the same number of reads (9M) for further downstream analysis. To calculate the enrichment, we used the BEDtools software suite (v.2.25.0) [64] to obtain the number of reads overlapping regions of interest. Resulting read counts were normalized (RPKM), input corrected and log2 transformed using R. We conducted peak calling using the MACS2 (v.2.1.1) [30] “callpeak” module with -t and -c options and default parameters, except for -g 2.73e8 –keep-dup all -B –SPMR -q 0.01 –nomodel. We further assessed statistical significance of MACS2 peaks using the RECAP software [65] to recalibrate peak calling P-values. Over 99% of MACS2 peaks remained significant according to a recalibrated P-value threshold of 0.05 (Table S2). For visualization in IGV [66, 67], tracks of input corrected ChIP-seq signal were computed using the MACS2 “bdgcmp" module (-m ppois) on each pair of fragment pileup and control lambda bedGraph files from peak calling analysis (https://github.com/taoliu/MACS/wiki/Build-Signal-Track).

To quantitatively compare histone modification profiles between infected and control mosquito tissues, we used diffReps software [33]. This method uses a sliding window approach to identify regions that show significant changes in ChIP-seq signal, without constraining regions to compare by peak calling. ChIP-seq data for infected and control and the corresponding inputs were provided with -tr, -co, –btr and –bco options. Due to the lack of biological replicates, the statistical test used was G-test (-me gt). Other parameters were set as default except for –window 1000, as recommended for the scanning of histone modification peaks. We performed annotation of MACS2 peaks and diffReps regions to genomic features (TSSs, exons, introns, and intergenic regions) using the annotatePeaks.pl module in HOMER (v.3.12) [68]. Based on the density distribution of the distances from upstream MACS2 peaks and diffReps regions to the nearest ATG site, we considered 2 Kb upstream from the translation start site ATG as the putative promoter region.

We used the chromatin-state segmentation software ChromHMM [31] to compute genome-wide chromatin state predictions in each condition based on relative enrichment levels of histone modifications. For the binarization of the genome we used default parameters except for -b 200. We chose a 4 state model assuming chromatin states with high levels of enrichment of each histone modification. The chromatin states we found were: depleted (low levels of enrichment for all histone modifications, high levels of enrichment for H3K9me3, high levels of enrichment for H3K9me3 and H3K4me3 (bivalent state), and high levels of enrichment for H3K9ac, H3K27ac and H3K4me3. According to the ChromHMM segmentation, most of the genome is in a depleted state. In Drosophila H3K27me3 occupies regions where H3K27ac is absent [47]. However, this histone modification mark is not included in our study. We assigned predicted chromatin states to different features, such as MACS2 peaks and diffReps regions, using the intersect tool from BEDtools, and we required a minimum overlap between the regions of 51% (-f 0.51).

To obtain a high confidence set of diffReps regions we applied a filtering based on multiple thresholds. We filtered out those diffReps regions located in depleted chromatin states (ChromHMM segmentation) at each corresponding condition and displaying FDR > 0.05. We also divided regions in three quantile groups (cut2 function in the Hmisc R package) according to their mean values in log2 Fold Change and in average normalized counts and fold enrichment vs input at each corresponding condition. Regions that fall in the lower quantiles were discarded (Table S3).

We performed GO terms overrepresentation tests for set of genes of interest annotated to diffReps regions using PANTHER Overrepresentation Test [69, 70]. We chose Fisher’s Exact with FDR multiple tests correction and applied a threshold of FDR < 0.05. Sets of differentially expressed genes between conditions, infected and control, were annotated based on the Gene Ontology (GO) and Kyoto Encyclopedia of Genes and Genomes (KEGG) using DAVID. We used ChromHMM segmentation and plotEnrichment function in chromstaR R package [71] to assess enrichment of predicted chromatin states in certain features of interest, e.g subset of genes encoding immune response factors. Average profile plots and heatmaps representing enrichment of histone modifications (RPKM normalized and input-corrected) centered on gene coordinates were built using ngs.plot (v.2.61) [72].

### Integration of RNA and ChIP-seq data

To connect patterns of histone modifications with the regulation of gene expression we ordered genes annotated to MACS2 peaks by mRNA levels and showed histone modification enrichment levels at those gene bodies and promoter regions. Correlation between histone modification enrichment levels and mRNA levels was assessed using Spearman’s rank correlation test (cor.test R function). Additionally, in order to connect differential enrichment of histone modifications with regulation of gene expression, we filtered those genes containing high-confidence diffReps regions in promoters and gene bodies and performed a soft clustering approach using the Mfuzz R package [36] over the ratio of histone modifications between conditions (ratio of enrichment at infected to uninfected samples). Using a standard m fuzzy c-means parameter of 1.7, a total of 30 clusters were created. Clusters with certain patterns of histone modifications were isolated creating unique groups (Figure S4). Only elements with membership value higher than 51% within each particular cluster were considered. Next, we used the clustering order based on the ratio of histone modification enrichment to show mRNA levels of the corresponding genes (ratio of mRNA levels at infected to un-infected samples). To check the validity of our results and to further assess the functional output and transcriptional shift associated to different chromatin states we focused on patterns showing maximum enrichment of certain histone modification at both infected and uninfected. We categorized genes into high, medium or low expression groups at each condition by dividing the mRNA levels in three quantile groups according to their means (cut2 function) and we filtered out low-expressed genes. Based on the soft-clustering of each region and each gene mRNA level, we then isolated those cases where differential histone enrichment profiles, a gain/loss in active hPTMs or gain/loss in the repressive H3K9me3 modification, coincide with the expected functional output: up or down regulation of the gene. We also performed a soft clustering analysis with the same parameters as above computing histone modification enrichment ratios at promoters and gene bodies of significant differentially expressed genes according to DESeq2 (P-value < 0.05), with similar results (Figure S7-8, Table S6).

Heatmaps showing histone modification enrichment and mRNA levels were built using the iheatmapr R package [73]. Bar and violin plots were produced using the ggplot2 R package [74]. For comparative and visualization purposes, histone enrichment signals and mRNA levels were log2 transformed. When computing histone modifications enrichments and ratios, a pseudocount (0.1) was added to obtain finite values when input correcting (dividing by 0 in ratios) or converting the signal to log2 scale. When categorizing values in quantile groups according to their means, High, Medium and Low groups, we used the cut2 function in the Hmisc R package.

### Motif analysis

We conducted *de novo* motif analysis using HOMER software (v.3.12) [68] on the set of MACS2 peaks that intersect with high confidence diffReps regions and annotate to genes (promoters and gene bodies). For this analysis, we considered the center of the ChIP-seq peak region and slopped 100 bp in each direction. We limited the number of background sequences to double the number of ChIP-seq target sequences for each histone modification. Only motifs enriched in more than 5% of the target sequences and below a threshold P-value of 10E-15 were considered, and results corresponding to low complexity motifs and offsets or degenerate versions of highly enriched motifs were avoided. The purpose of this analysis was to identify enrichment in particular sequence footprints associated with changes in histone modification occupancy that are associated to *P. falciparum* infection.

## DATA AVAILABILITY

ChIP-seq and RNA-seq data are deposited in the GEO database under accession number GSE120076.

## ACKNOWLEDGEMENTS

We would like to thank all children and their parents who participated in this study, as well as the local authorities in Burkina Faso for their support. We are very grateful to B. Yameogo, S. Tamboula, R. Hien, J. Bazié., B. Dabiré., F. Da, A. Diasso and F. Yao in Burkina Faso for technical assistance. We thank The Genomic Services Lab at the HudsonAlpha Institute for Biotechnology, specially Braden Boone, Angela Jones, and Terri Pointer, for their help in preparing RNA-seq libraries and performing Illumina sequencing of RNA-seq and ChIP-Seq samples. The content is solely the responsibility of the authors and does not necessarily represent the official views of the National Institutes of Health.

## Author Contributions

E.G.-D. and V.G.C. conceived the study. E.G.-D., R.S.Y., T.L. and V.G.C. designed the experiments. E.G.-D. performed laboratory experiments. E.G.-D., R.S.Y. and T.L. performed the field experiments. J.L.R. and E.G.-D. performed data analysis. R.S.Y., T.L., and J.B.O. provided infrastructure and organised gametocyte carrier recruitment in Burkina Faso. E.G.-D., J.L.R. and V.G.C. wrote the manuscript. R.S.Y., and T.L. commented on the manuscript. All the authors revised the manuscript and approved the final version.

